# A Novel Oncogenic Function of PRC2 Heterogeneity in Medulloblastoma

**DOI:** 10.1101/2021.12.02.470979

**Authors:** Jiaqing Yi, Xuanming Shi, Xiaoming Zhan, Richard Q Lu, Zhenyu Xuan, Jiang Wu

## Abstract

Intratumor epigenetic heterogeneity is emerging as a key mechanism underlying tumor evolution and drug resistance. Medulloblastomas, the most common childhood malignant brain tumor, are classified into four subtypes including SHH medulloblastomas, which are characterized by elevated SHH signaling and a cerebellum granule neuron precursor (CGNP) cell-of-origin. Medulloblastomas are highly associated with epigenetic abnormalities. We observed that the histone H3K27 methyltransferase polycomb repressor complex 2 (PRC2) is often heterogeneous within individual SHH medulloblastoma tumors. Using mouse models, we showed that while a complete deletion of the PRC2 core subunit EED inhibited medulloblastoma growth, a mosaic deletion of EED significantly enhanced tumor growth. EED is intrinsically required for CGNP maintenance by inhibiting both neural differentiation and cell death. Complete EED deletion led to CGNP depletion and reduced occurrence of medulloblastoma. Surprisingly, we found that medulloblastomas with mosaic EED levels grew faster than did control wildtype tumors and expressed increased levels of oncogenes such as *Igf2. Igf2* is directly repressed by PRC2 and has been demonstrated to be both necessary and sufficient for SHH medulloblastoma progression. We showed that IGF2 mediated the oncogenic effects of PRC2 heterogeneity in tumor growth. Using a human medulloblastoma cell line, we generated clones with different EED levels and confirmed that EED^low^ cells could stimulate the growth of EED^high^ cells through derepressed IGF2 signals. Thus, PRC2 heterogeneity controls medulloblastoma growth through both intrinsic growth competence and non-cell autonomous mechanisms in distinct tumor subclones. We reveal a novel oncogenic function of PRC2 heterogeneity in tumor development.

## Introduction

Cancer development is driven by tumor-intrinsic mutations and is fueled by the tumor microenvironment (TME), which consists of various non-cancerous cells and cancer-derived subclones (1–4). Intratumor heterogeneity at the genetic or epigenetic levels is emerging as a widespread mechanism underlying tumor evolution and drug resistance (5, 6). An important question is how different cancer subclones compete and cooperate. Non-cell autonomous mechanisms affecting neighboring cancer cells could underlie the oncogenic function of tumor heterogeneity (7). The significance of cancer subclone cooperation has been demonstrated in several cancer types. For example, in glioblastomas, cancer cells expressing a mutant EGFR stimulate the growth of cancer cells with a wild-type EGFR through cytokine secretion (8). In pediatric glioblastoma, inactivating mutations in the histone methyltransferase *KMT5B*, present in□<1% of tumor cells, confer increased invasion and migration on neighboring cells through chemokine signaling(9). In small cell lung cancer, a fraction of cancer cells with activated NOTCH signaling could support the growth of NOTCH^low^ neuroendocrine cancer cells through a non-cell autonomous mechanism (10). Understanding the interactions between various cancer subclones could guide the development of new therapies.

Medulloblastomas, the most common childhood malignant brain tumor, are classified into four subgroups based on distinctive transcription profiles and epigenetic landscapes. These subgroups are referred to as WNT, SHH, Group 3, and Group 4 (11–15). Within these subgroups, SHH medulloblastomas are characterized by abnormally elevated Sonic hedgehog (SHH) signaling in cerebellum granule neuron precursors (CGNPs). During early postnatal development, CGNPs expand in a SHH-dependent manner and differentiate to granule neurons. A balanced CGNP proliferation and differentiation is precisely controlled by SHH signaling and additional factors. Abnormally elevated SHH signaling caused by various mutations leads to CGNP overexpansion and eventually medulloblastoma. SHH medulloblastomas are highly heterogeneous at multiple levels including cytopathology, patient age, prognosis, and genomic and epigenetic abnormalities (16, 17). The inter-tumor heterogeneity is believed to be caused by oncogenic pathways in addition to SHH signaling (16, 18). Recent single-cell RNA-seq (scRNA-seq) studies further revealed intratumor heterogeneity in SHH medulloblastomas (19–21). SHH medulloblastomas contain mainly ATOH1^+^ CGNP-like tumor propagating cells and NEUROD1^+^ differentiated granule neurons. Other minor cell populations include OLIG2^+^ cancer stem cells (19), other neuron subtypes, astrocytes, microglia (22), fibroblasts, and blood vessel cells. We and others also observed the recruitment of immune cells to SHH medulloblastomas (23, 24). These studies demonstrated the complexity of SHH medulloblastomas and the sensitivity of SHH medulloblastomas to TME changes. Within single human medulloblastoma tumors, subclones that harbor different genetic mutations and recurrent potentials have been observed (25). However, it is not clear how these tumor subclones interact with each other and affect overall tumor growth.

Epigenetic regulators often play context-dependent roles in cancer development through both oncogenic and tumor suppressor functions. This is likely through the regulation of specific sets of target genes in different cell types and at specific tumor developmental stages (26–30). One intriguing discovery from genomic studies of medulloblastoma is that there is a high rate of alterations of epigenetic regulators across all medulloblastoma subgroups. More than 30% of medulloblastomas harbor mutations, deletions, or amplifications of genes that encode histone modification enzymes or chromatin remodelers. Each medulloblastoma subtype also displays common and distinct epigenetic properties (31–33). These findings suggest that epigenetic alterations are important drivers of medulloblastoma. One dysregulated epigenetic pathway identified in medulloblastoma involves histone H3 lysine 27 trimethylation (H3K27me3) (32, 34). The methyltransferase PRC2 adds the repressive mark H3K27me3, and thus is a major transcription repressor. In many cancers, PRC2 displays oncogenic functions by repressing tumor suppressors or differentiation processes (35–38), but PRC2 can also act as a tumor suppressor by repressing oncogenes (36, 39–41). The PRC2 core subunits EZH2/EZH1, EED, and SUZ12 are highly expressed in medulloblastoma with no loss-of-function mutations identified (32). It has been speculated that PRC2 is oncogenic in medulloblastoma (15). PRC2 inhibitors have been used to inhibit medulloblastoma growth in xenograft models (42, 43). However, a recent genetic study of Group 3 medulloblastoma mouse models showed an unexpected tumor-suppressor function of PRC2 (44). In addition, a study of PRC2 antibody staining in 88 medulloblastoma tumor specimens showed that EZH2+ cell numbers vary from 20% to 80% among different tumors. The prognosis outcome is not simply associated with a higher or lower percentage of EZH2+ cells (45). Thus, PRC2 may play complex roles in medulloblastoma development.

In this study, we identified a novel oncogenic function of PRC2 heterogeneity in medulloblastoma development. We observed that PRC2 levels are often heterogeneous within individual SHH medulloblastoma tumors. Using mouse models, we showed that while a complete deletion of the PRC2 core subunit EED inhibited medulloblastoma growth, a mosaic deletion of EED significantly enhanced tumor growth. Complete EED deletion leads to CGNP depletion and a reduced occurrence of medulloblastoma. Surprisingly, we found that medulloblastoma with mosaic EED levels grew faster than control wildtype tumors. We showed that IGF2 that was derepressed from partial EED deletion mediated the oncogenic effects of PRC2 heterogeneity in tumor growth. Using a human medulloblastoma cell line with different EED levels, we confirmed that EED^low^ cells could stimulate the growth of EED^high^ cells via the IGF2/PI3K/AKT signaling pathway. Thus, PRC2 heterogeneity controls medulloblastoma growth through both intrinsic growth competence and non-cell autonomous mechanisms in distinct tumor subclones.

## Results

### EED is required for normal mouse cerebellar development

SHH medulloblastoma is caused by abnormally active SHH signaling in CGNP cells. To determine the function of PRC2 in CGNPs and medulloblastomas, we deleted the PRC2 core subunit *Eed* specifically in CGNPs using a knock-in pan CGNP *Atoh1-Cre* driver (46) (Figure S1A). *Eed* deletion results in the degradation of other PRC2 subunits and loss of PRC2 function (47). Atoh1-Cre-mediated *Eed* deletion led to severe ataxia and reduced body weight in mutant mice (Figure 1A). In mature cerebella at postnatal day 28 (P28), H&E and NeuN staining showed that *Eed* deletion caused close-to-complete depletion of granule neurons without significantly altering the cerebellar foliation (Figure 1B). In *Eed*-deleted cerebella during development at P5, EED protein deletion and a decrease in H3K27me3 were prominent in most CGNPs (Figure 1C). We observed a gradual decrease in thickness of the external granule layer (EGL), where CGNPs reside, from P2 to P7, indicating an early depletion of CGNPs (Figure S1B).

**Figure 1.**
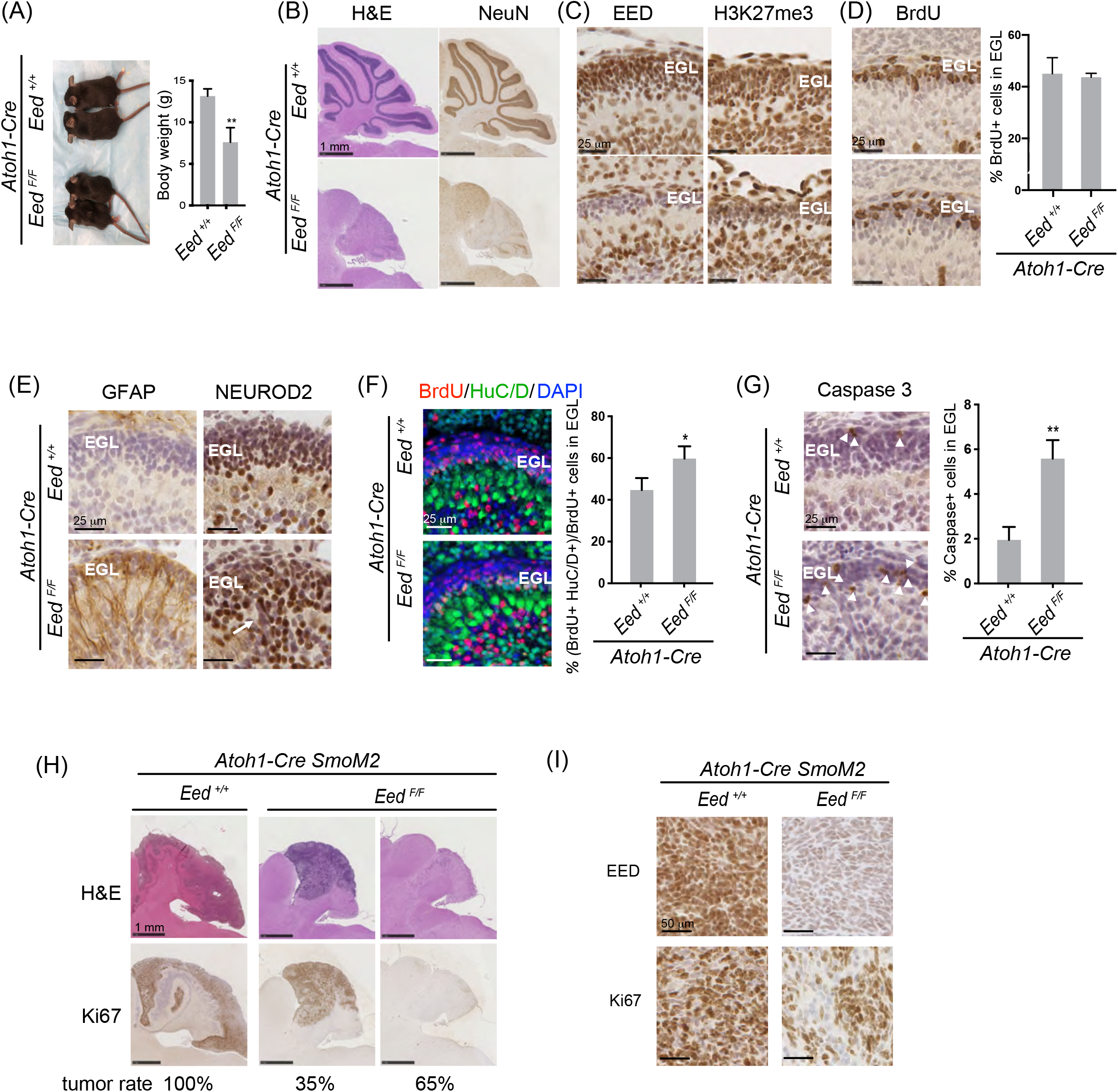
EED is required for cerebellar development and medulloblastoma formation. (A) Picture shown was *Atoh1-Cre Eed^+/+^* and *Atoh1-Cre Eed^F/F^* mice at P28. Quantification of body weight is shown on the right (n=4). (B-G) H&E and immunostaining of sagittal sections of cerebella from *Atoh1-Cre Eed^+/+^* and *Atoh1-Cre Eed^F/F^* mice at P28 (B), P5 (C, E G), and P2 (D, F). BrdU labeling was 30 min in (D) and 24 h in (F). Quantifications are shown on the right (n=3). (H-I) H&E and immunostaining of sagittal sections of cerebella from P28 *Atoh1-Cre SmoM2 Eed^+/+^* and *Atoh1-Cre SmoM2 Eed^F/F^* mice. Percentages under each genotype in (H) indicates their respective tumor occurrence rate (n=20 per group). Student’s t-test, *, p<0.05; **, p<0.01. **Please also see Figure S1**.

Previous studies demonstrated cell type and developmental stage specific functions of the PRC2 complex in controlling the timing of progenitor proliferation and neuronal/glial differentiation (48–50). Using BrdU pulse labeling (30 min), we did not observe a significant difference in the percentage of BrdU+ cells in control and *Eed*-deleted EGLs (Figure 1D), suggesting that EED does not affect CGNP proliferation rate. On the other hand, we observed a significant increase in GFAP staining in the *Eed*-deleted P5 cerebellum (Figure 1E). Since CGNP normally is restricted to the granule neuron fate, EED deletion may result in an abnormal fate change to glial cells. We previously showed that a proneural gene *NeuroD2* is activated by H3K27me3 demethylase UTX and promotes granule neuron differentiation (24). Antibody staining showed that NEUROD2 was expressed in differentiating granule neurons in the inner EGL and mature neurons in the inner granule layer (IGL) in the control P5 cerebellum but displayed increased expression and abnormal distributions in the *Eed*-deleted cerebellum (Figure 1E, arrow). In a BrdU tracing experiment, in which pups were injected with BrdU 24 h before examination, the newly differentiated neurons were co-labeled with BrdU and a neuronal marker HuC/D. The percentage of BrdU+ HuC/D+ cells within all BrdU+ cells was significantly higher in *Eed*-deleted EGLs (Figure 1F), confirming an enhanced neuronal differentiation upon EED deletion. Moreover, increased active caspase 3 staining showed an increase of cell death in *Eed*-deleted EGLs than in control EGLs (Figure 1G, arrow heads). Together, these results indicate that EED is required for normal neuron/glial differentiation, migration, and survival in the cerebellum. EED deletion led to close-to-complete CGNP and granule neuron depletion.

### EED is required for SHH medulloblastoma formation

In a well-characterized SHH medulloblastoma mouse model, which was caused by a Cre-induced constitutively active *SmoM2* mutation (51), we deleted *Eed* from precancerous CGNPs using *Atoh1-Cre*. Deletion of *Eed* from medulloblastoma mice also caused CGNP depletion and severe ataxia. As a result, compared to the 100% medulloblastoma occurrence in *SmoM2* mice with wild-type *Eed*, EED deletion reduced cancer occurrence rate to about 35%, whereas 65% of the *Eed*-deleted mice were tumor free (Figure 1H). The *Eed*-deleted tumors were completely EED negative and showed lower proliferation rates by Ki67 staining (Figure 1I). Therefore, consistent with its oncogenic role in many cancers, the PRC2 complex is required for progenitor and tumor development in SHH medulloblastoma, likely in a cell-autonomous manner.

### Heterogeneous TgAtoh1-Cre activity led to incomplete EED deletion and mild cerebellar defects

Using another well-characterized CGNP-specific transgenic *Atoh1-Cre* line (*TgAtoh1-Cre*) (52), we observed an unexpectedly mild phenotype in *Eed*-mutant cerebella. *TgAtoh1-Cre Eed^F/F^* mice were viable and did not show obvious motor defects. Anatomically, we only observed minor changes in mutant cerebella at P28, in which the granule neurons in the IGL were less dense than those in control cerebella (Figure 2A). This defect in granule neuron loss was more obvious in the anterior lobes than in the posterior lobes. In developing cerebella at P8, we observed various levels of EED deletion in the *TgAtoh1-Cre Eed^F/F^* EGL as shown by antibody staining. More EED-deleted CGNPs were observed in anterior lobes than in posterior regions (Figure 2B). Similar to the EED-deleted *Atoh1-Cre Eed^F/F^* cerebella, the anterior EGL regions of *TgAtoh1-Cre Eed^F/F^* mice at P5 displayed increased GFAP expression and increased caspase 3 staining than control mice (Figure 2C), confirming a function of EED in preventing abnormal differentiation and cell death.

**Figure 2.**
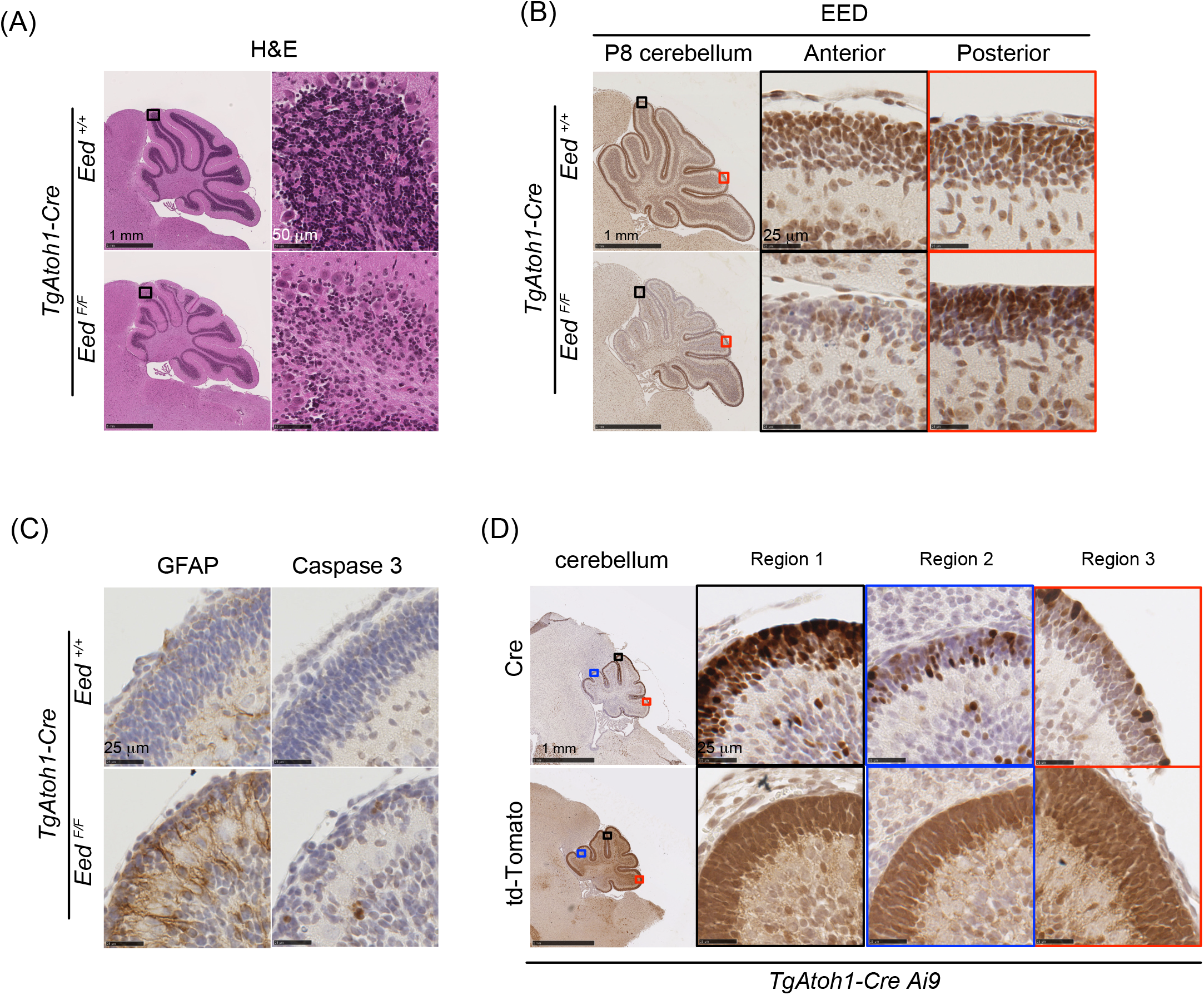
Heterogeneous TgAtoh1-Cre activity led to incomplete EED deletion and mild cerebellar defects. (A) H&E staining of sagittal sections of cerebella from P28 *TgAtoh1-Cre Eed^+/+^* and *TgAtoh1-Cre Eed^F/F^* mice. The boxed areas in the left pictures are shown on the right. (B-C) Immunostaining of sagittal sections of cerebella from P7 *TgAtoh1-Cre Eed^+/+^* and *TgAtoh1-Cre Eed^F/F^* mice. Pictures on the right in (B) are boxed areas in the pictures on the left with corresponding outline colors. (D) Cre and td-Tomato staining of sagittal sections of cerebella from P2 *Ai9 TgAtoh1-Cre* mice. Pictures on the right are boxed regions in the pictures on the left with corresponding outline colors. **Please also see Figure S2**.

The incomplete deletion of EED led us to re-examine the TgAtoh1-Cre expression and activities. Although high levels of Cre proteins were detected throughout the EGLs in P5 cerebella, their levels were heterogeneous. Cre levels were relatively high in some anterior EGL regions, but were more mosaic in other regions (Figure 2D). The Cre activities were examined using the *Ai9* reporter line that contains an *R26R-floxed stop-tdTomato* allele (53). Interestingly, TgAtoh1-Cre-mediated recombination led to the expression of tdTomato in almost all CGNPs in EGL at P5 (Figure 2D), which is consistent with previous observations that the *TgAhoh1-Cre* allele is widely active in CGNPs (52). Together, these results indicate that there were heterogeneous levels of Cre proteins expressed in *TgAtoh1-Cre* CGNPs, which we termed Cre^high^ and Cre^low^ CGNPs. In *TgAtoh1-Cre Ai9* mice, the Cre activities in both Cre^high^ and Cre^low^ CGNPs were sufficient to recombine the “easy” *R26R-floxed stop-tdTomato* locus. However, in *TgAtoh1-Cre Eed^F/F^* mice, while Cre^high^ CGNPs could delete EED, low Cre activities in Cre^low^ CGNPs failed to completely recombine the more “difficult” *Eed^F^* allele, resulting in a mosaic deletion of EED in CGNPs and mild cerebellum defects.

Besides the Cre activities, the other difference between the *Atoh1-Cre* and *TgAtoh1-Cre* is that the knock-in *Atoh1-Cre* led to *Atoh1* heterozygosity, whereas *TgAtoh1-Cre* mice has two wildtype *Atoh1* copies. To determine whether *Atoh1* dosage could contribute to the different patterns of EED deletion in these two Cre mice, we generated the *TgAtoh1-Cre Atoh1^+/-^ Eed^F/F^* mice (Figure S2A). Although losing one copy of *Atoh1* from the *TgAtoh1-Cre Eed^F/F^* mice led to a more severe loss of granule neurons in P28 cerebella, many neurons in the posterior lobes remain (Figure S2B). EED staining also showed a mosaic EED deletion pattern in *TgAtoh1-Cre Atoh1^+/-^ Eed^F/F^* CGNPs in P8 EGLs. Thus, the incomplete EED deletion in *TgAtoh1-Cre Eed^F/F^* CGNPs was mostly caused by the heterogeneous *TgAtoh1-Cre* activities.

### *TgAtoh1-Cre SmoM2 Eed^F/F^* mice showed enhanced tumor growth with mosaic EED deletion

Strikingly, in contrast to the depletion of CGNPs and reduced tumor occurrence after complete EED deletion in the *Atoh1-Cre SmoM2 Eed^F/F^* mice, the *TgAtoh1-Cre SmoM2 Eed^F/F^* mice had enhanced tumor growth and shortened survival time compared to control wildtype and *Eed^F/+^* tumor mice. Compared to a 38-day median survival time for *TgAtoh1-Cre SmoM2 Eed^+/+^* mice, *TgAtoh1-Cre SmoM2 Eed^F/F^* mice had only a 23-day median survival time with aggressive tumor growth (Figure 3A, 3B). Compared to wildtype tumors, *Eed^F/F^* tumors had increased proliferation rates as shown by BrdU labeling (Figure 3C) and decreased cell death rate as shown by active caspase 3 staining (Figure 3D). Deleting one copy of *Atoh1* from the *TgAtoh1-Cre SmoM2* mice delayed the tumor occurrence and progression for all three *Eed* genotypes. The survival times were still significantly shortened in *Eed^F/F^* mice compared to *Eed^+/+^* and *Eed^F/+^* mice (Figure S2D). These results suggest an unexpected tumor suppressor function of PRC2 in SHH medulloblastoma.

**Figure 3.**
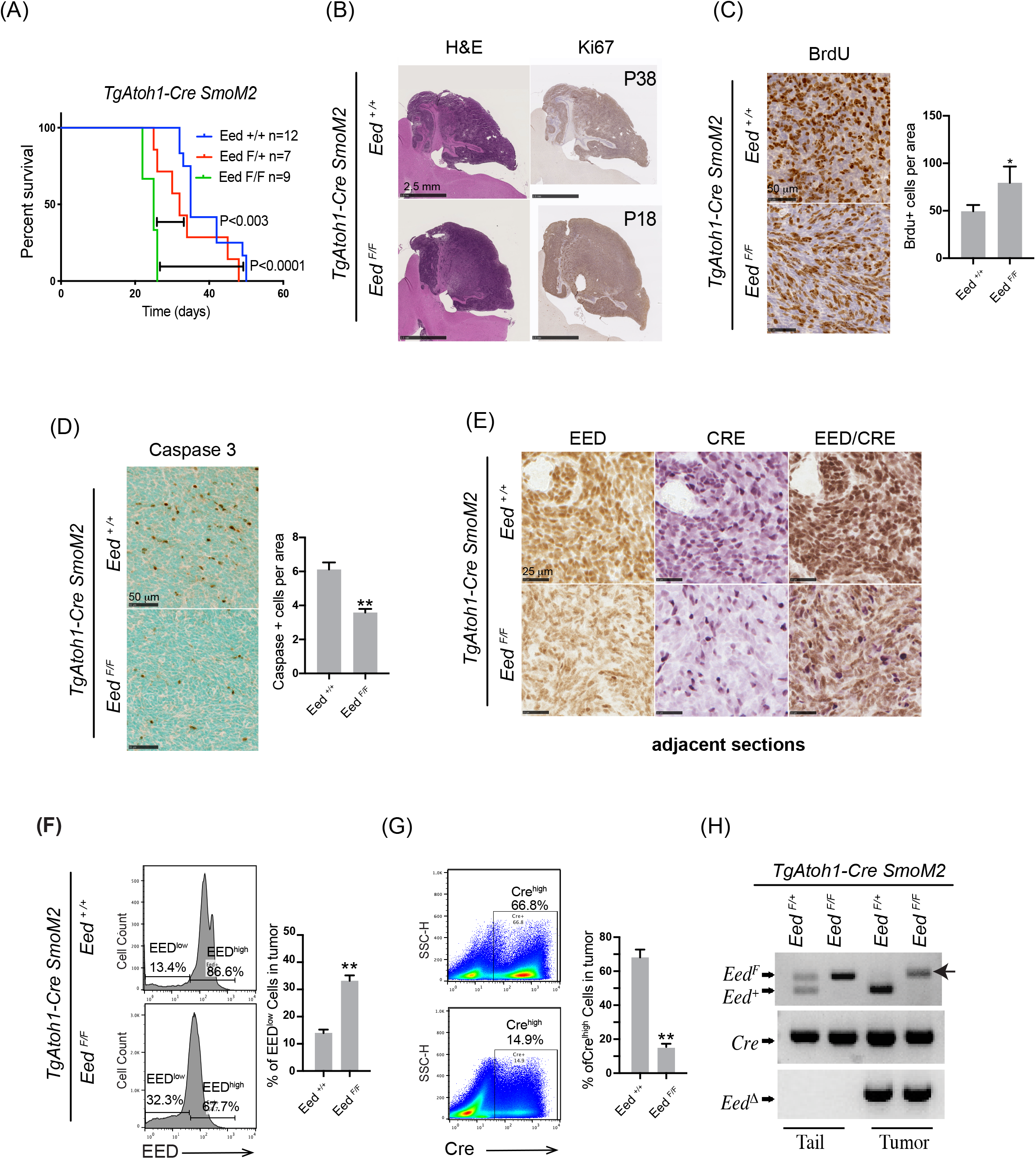
*TgAtoh1-Cre SmoM2 Eed^F/F^* mice showed enhanced tumor growth with mosaic EED deletion. (A) Survival curves of mice harboring *TgAtoh1-re SmoM2* medulloblastoma with indicated *Eed* genotypes. (B) H&E and Ki67 staining of sagittal sections of tumor from *TgAtoh1-Cre SmoM2 Eed^+/+^* mice at P38 and *TgAtoh1-Cre SmoM2 Eed^F/F^* mice at P18. (C-E) Immunostaining of sagittal sections of tumor from P18 *TgAtoh1-Cre SmoM2 Eed^+/+^* and *TgAtoh1-Cre SmoM2 Eed^F/F^* mice. Quantification is shown on the right (n=4). BrdU labeling time was 30 min in (C). (F-G) Flow cytometric analyses of EED (F, histograms, n=3) and Cre (G, dot plots, n=3) distribution in tumors from P18 *TgAtoh1-Cre SmoM2 Eed^+/+^* and *TgAtoh1-Cre SmoM2 Eed^F/F^* mice. (H) PCR genotyping of tail and tumor tissues from *TgAtoh1-Cre SmoM2 Eed^F/+^* and *TgAtoh1-Cre SmoM2 Eed^F/F^* mice. The arrow points to the unrecombined *Eed^F^* allele in the *TgAtoh1-Cre SmoM2 Eed^F/F^* tumor. Student’s t-test, *, p<0.05; ****, p<0.0001. **Please also see Figure S2**.

We then examined EED and Cre protein expression in the *TgAtoh1-Cre SmoM2* tumors. In *Eed^+/+^* wildtype tumors, both EED levels and Cre levels were high in most tumor cells, as expected. There were also some variations in EED and Cre levels, which reflects the different cell types in the tumor as well as the heterogeneous TgAtoh1-Cre expression pattern (Figure 3E). A more quantitative measurement using FACS showed that in wildtype medulloblastoma tumors, more than 85% of tumor cells were EED^high^ and more than 65% were Cre^high^ cells (Figure 3F, 3G). Interestingly, in *Eed^F/F^* mutant medulloblastomas, many cells still expressed EED (Figure 3E). FACS analyses showed that in *Eed^F/F^* medulloblastomas, ~68% of the cells retained EED. The EED^low^ cells were increased in *Eed^F/F^* tumors than in *Eed^+/+^* tumors, but they only consisted of ~32% of all *Eed^F/F^* tumor cells (Figure 3F). Intriguingly, in *Eed^F/F^* medulloblastomas, Cre^high^ cells were significantly reduced to ~15% (Figure 3E, 3G). Therefore, in *TgAtoh1-Cre* induced medulloblastomas, Cre^low^ cells are likely a minor population but are disproportionally expanded in *Eed^F/F^* mutant medulloblastomas, contributing to overall faster tumor growth. The Cre^low^ population likely retained at least one wildtype *Eed* allele, as confirmed by co-staining and genotyping (Figure 3E, 3H). In these *Eed^F/F^* mutant medulloblastoma, there are subclones of tumor cells with different Cre activities and EED levels (Cre^high^ EED^low^ and Cre^low^ EED^high^). The EED deletion in Cre^high^ EED^low^ subclones may lead to the overexpansion of the Cre^low^ EED^high^ population and faster overall tumor growth. During tumor progression, the growth competition pressure would selectively maintain a high percentage of EED^high^ cells and a low percentage of EED^low^ cells.

### *Igf2* is a PRC2 target that was derepressed in *TgAtoh1-Cre SmoM2 Eed^F/F^* tumors

To identify EED-regulated genes and pathways involved in promoting mosaic medulloblastoma growth, we performed RNA-seq and compared transcriptomes of the *TgAtoh1-Cre SmoM2 Eed^+/+^* medulloblastomas with those of the mosaic *TgAtoh1-Cre SmoM2 Eed^F/F^* tumors (Figure 4A). The results showed that few genes involved in the core SHH pathway or in the neural developmental process were changed significantly (Figure S3). The level of key SHH medulloblastoma signature genes such as *Gli1, Atoh1*, and *NeuroD1* were similar between *Eed^+/+^* and *Eed^F/F^* tumors, indicating that the fast growth of *Eed^F/F^* medulloblastomas was likely not due to changes in the tumor cell-intrinsic fate. The cancer stem cell markers *Olig2* and *CD133*, as well as the recently identified glia/microglia/Igf1 TME pathway genes (22), were also not changed (Figure S3). Of the 231 upregulated genes in the *Eed^F/F^* mutant medulloblastoma (fold change >2, p<0.05), many are classical PRC2 targets such as *Hox* gene clusters, muscle development genes, and tumor suppressors such as *Cdkn1c* (Figure 4A, 4B, S3). Notably, some of the upregulated genes in *Eed^F/F^* mutant medulloblastomas encode oncogenic molecules such as IGF2, NOTCH1, and LIN28b (54, 55) (Figure 4A-B). Using RT-qPCR, we confirmed that *Igf2* was increased in *TgAtoh1-Cre SmoM2* medulloblastoma when *Eed* was deleted (Figure 4C). We performed H3K27me3 ChIP-seq of *SmoM2* medulloblastoma and observed extensive H3K27me3 signals around the EED-repressed oncogenes including *Igf2* (Figure 4D). As an imprinting gene, *Igf2* has been shown to be repressed by the PRC2 complex in other systems (56, 57). Using ChIP-qPCR, we showed that EED directly binds to the *Igf2* promoter (Figure 4E). Upon EED deletion in the *Eed^F/F^* medulloblastomas, EED binding as well as local H3K27me3 levels were reduced (Figure 4F). The main signaling pathway downstream of IGF2 is the PI3K/AKT pathway, and high levels of phosphorylated AKT (p-AKT) were observed in some SHH medulloblastoma patients with poor prognoses (16). Indeed, we observed increased p-AKT levels in *Eed^F/F^* mutant medulloblastomas compared to wild-type medulloblastomas (Figure 4G). Thus, *Igf2* is a PRC2 target that was derepressed in *TgAtoh1-Cre SmoM2 Eed^F/F^* tumors.

**Figure 4.**
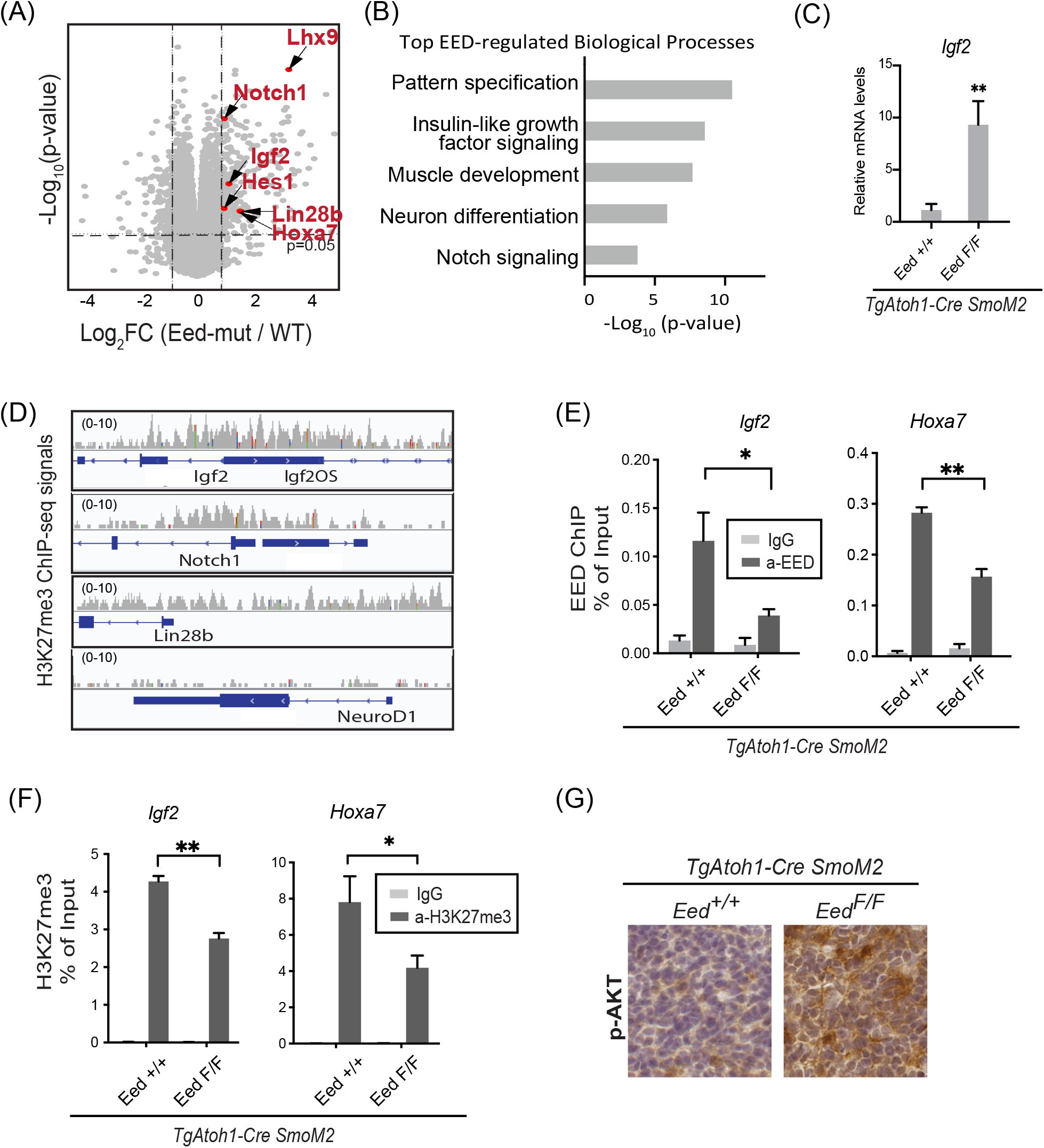
*Igf2* is a PRC2 target that is derepressed in *TgAtoh1-Cre SmoM2 Eed^F/F^* tumor. (A) Volcano plot of RNA-seq data comparing *TgAtoh1-Cre SmoM2 Eed^+/+^* (WT) and *TgAtoh1-Cre SmoM2 Eed^F/F^* (*Eed-mut*) tumors. Red-labelled dots are known oncogenes increased in *Eed-mut* tumors. (B) Gene ontology analysis of enriched pathways in upregulated genes in *EED-mut* tumors. (C) RT-qPCR of *Igf2* expression in *TgAtoh1-Cre SmoM2 Eed^+/+^* and *TgAtoh1-Cre SmoM2 Eed^F/F^* mice (n=6). (D) H3K27me3 ChIP-seq signals for *Igf2, Notch1*, and *Lin28b. NeuroD1* was used as an H3K27me3 negative control. (E-F) ChIP-qPCR analysis of EED (E) and H3K27me3 (F) at the promoter regions of *Igf2* and *Hoxa7* (n=3) in *TgAtoh1-Cre SmoM2 Eed^+/+^* and *TgAtoh1-Cre SmoM2 Eed^F/F^* tumors. (G) p-AKT staining of sagittal sections of tumors from *TgAtoh1-Cre SmoM2 Eed^+/+^* and *TgAtoh1-Cre SmoM2 Eed^F/F^* mice. Student’s t-test, *, p<0.05; **, p<0.01. **Please also see Figure S3**.

### Second mouse model demonstrating the oncogenic function of PRC2 heterogeneity

Our genetic experiments suggest that a mosaic deletion of EED led to faster tumor growth, possibly through derepressed IGF2 signaling. To exclude the possibility of unknown effects of the expression of TgAtoh1-Cre, we induced mosaic EED deletion in SHH medulloblastoma in another mouse model. Previously, we developed a genetic system able to delete specific genes in medulloblastoma after tumor formation (28). *CAG-CreER SmoM2 Eed^F/F^* mice develop SHH medulloblastoma spontaneously (51). Basal Cre activities could induce *SmoM2* at the *Rosa26* locus but are not sufficient to delete *Eed. Eed* could then be deleted by tamoxifen induction in a dose-dependent manner. We transplanted the *CAG-CreER SmoM2 Eed^F/F^* medulloblastomas subcutaneously to NOD/SCID mice, which were then injected with tamoxifen or an oil control (Figure 5A). The control transplanted tumor cells with oil injection were EED^high^ (Figure 5B, 5C). When tamoxifen was injected 10 times (10xTAM) post transplantation, EED was mostly deleted from the tumor cells, and tumor growth was significantly inhibited (Figure 5B, 5C). On the contrary, injecting tamoxifen only once (1XTAM), which partially deleted EED in transplanted tumors, resulted in enhanced tumor growth (Figure 5B, 5C). In EED heterogeneous tumors, *Igf2* expression level was increased compared to the oil-treated homogeneous EED^high^ tumors (Figure 5D). Thus, using a second genetic system, we confirmed that while a complete EED deletion led to tumor inhibition, a mosaic EED deletion promoted medulloblastoma growth, possibly through the increased IGF2 signaling.

**Figure 5.**
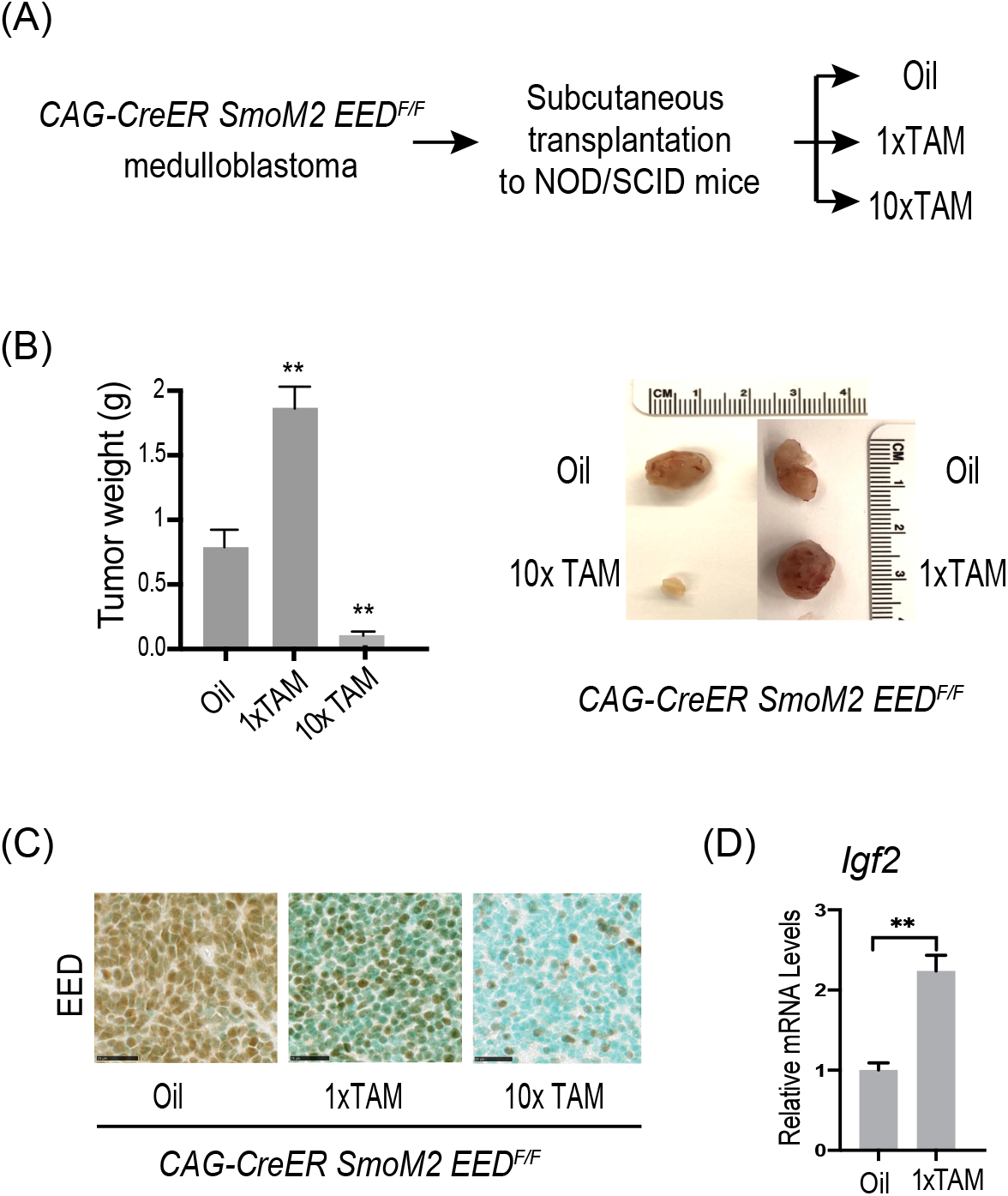
Second mouse model demonstrating the oncogenic function of PRC2 heterogeneity. (A) Experimental scheme of deleting EED completely or mosaically after SHH medulloblastoma tumor formation. (B) Comparison of the growth of *CAG-CreER SmoM2 Eed^F/F^* medulloblastoma transplanted subcutaneously to NOD/SCID mice treated with either oil, one or 10 doses of TAM (n=3). Representative images of transplanted tumors are shown on the right. (C) EED staining of sections of transplanted tumors from NOD/SCID mice treated with either oil, one or 10 doses of TAM. (D) RT-qPCR of *Igf2* expression comparing tumors from NOD/SCID mice treated with either oil or one dose of TAM (n=3).

### PRC2 heterogeneity and IGF2 expression in human medulloblastoma samples

*IGF2* is especially of interest to SHH medulloblastoma research because it is specifically expressed at high levels in human SHH medulloblastomas compared to other medulloblastoma types, based on the analysis of a cohort of 172 medulloblastoma patients (58) (Figure 6A). Previous genetic studies also showed that IGF2 is both necessary and sufficient for promoting SHH medulloblastoma progression in mice (54, 59). Our results suggest that the *Igf2* gene is directly repressed by PRC2 in SHH medulloblastoma. Thus, IGF2 could be a key signaling molecule that is derepressed in EED^low^ cells and stimulates the growth of EED^high^ cells. After analyzing a previously published SHH medulloblastoma microarray data set (18), we also observed an overall increase in *IGF2* expression levels in SHH medulloblastomas with lower *EED* levels compared to the group with higher *EED* levels (Figure 6B).

**Figure 6.**
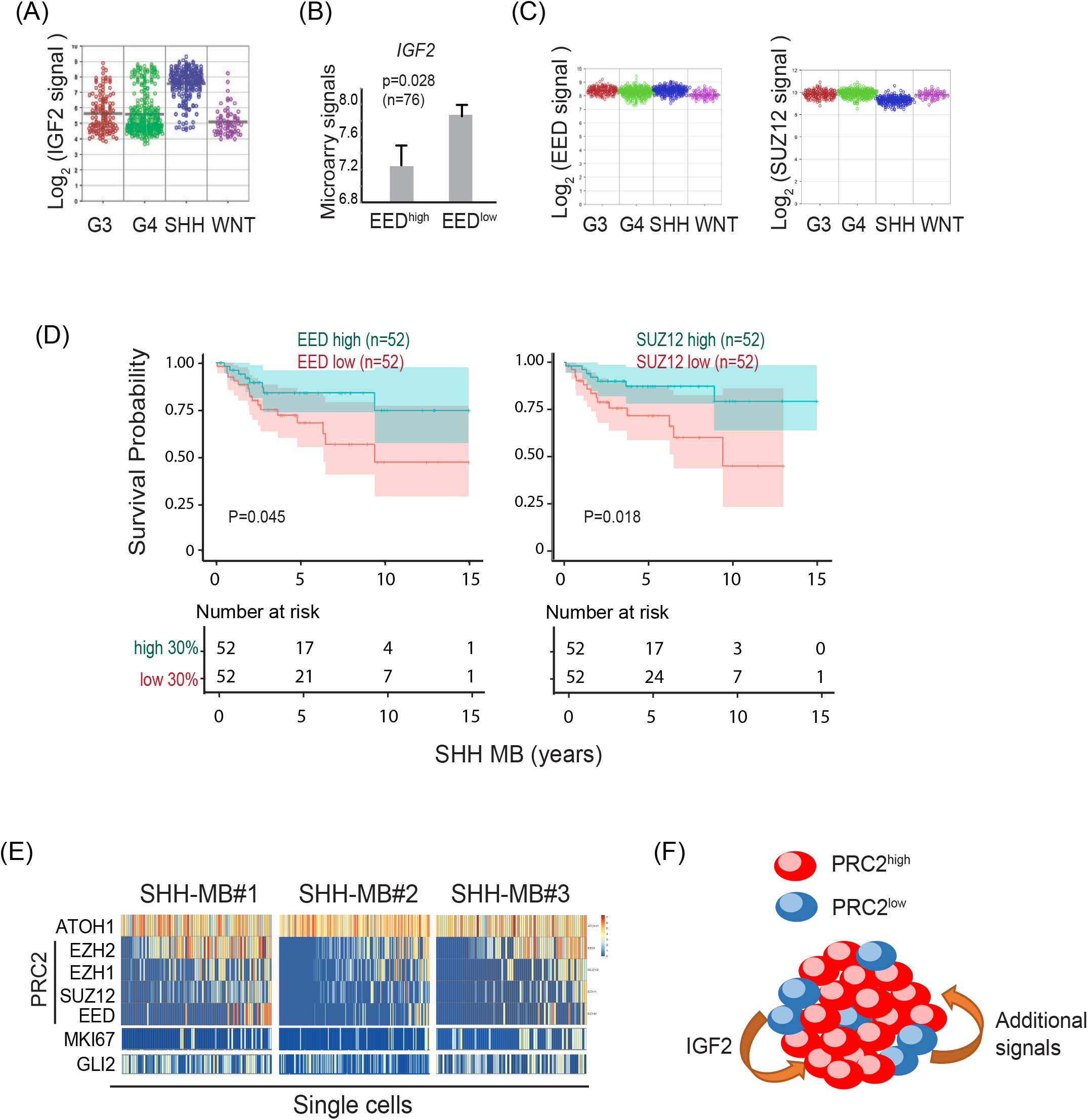
PRC2 heterogeneity and IGF2 expression in human medulloblastoma samples. (A) RNA-seq signals of *IGF2* among different subgroups of human medulloblastoma (n=172). (B) *IGF2* levels are inversely related to *EED* in human SHH medulloblastoma. Shown are average microarray signals of *IGF2* in SHH medulloblastomas with high or low EED levels (n=76) (16). (C) RNA-seq signals of *EED* and *SUZ12* among different subgroups of human medulloblastoma (n=172). (D) Kaplan-Meier curves with 95% confidence interval of human SHH medulloblastoma with different levels of *EED* expression (left) and *SUZ12* (right) (n=172). Shown are the comparisons between the top 30% and bottom 30% of patients ranked by *EED* or *SUZ12* tumor expression. (E) Heatmap showing the expression of PRC2 subunits, *MKI67* and *GLI2* in all the *ATOH1^+^* cells in the Smart-seq of the 3 human SHH medulloblastoma samples (20). Each column represents a single cell and the columns are arranged according to the sum of PRC2 subunit levels. Blue indicates undetectable levels. (F) The medulloblastoma subclone cooperation model. In a heterogeneous tumor that contains both PRC2^high^ and PRC2^low^ cells, the PRC2l^ow^ subclones, despite a lower growth competence, enhance PRC2^high^ subclone growth through derepressed secreted growth factors such as IGF2.

PRC2 subunits are highly expressed in all 4 medulloblastoma subgroups (Figure 6C). When separated by tumor PRC2 subunit expression levels, a prognosis analysis of the SHH medulloblastoma patients showed that relatively low levels of both *EED* and *SUZ12* correlated with worse prognosis outcomes (Figure 6D). These results suggest that PRC2 has a tumor suppressor function in addition to its potential oncogenic function in SHH medulloblastoma. To further determine whether PRC2 heterogeneity could underly the seemingly paradoxical functions of PRC2 in SHH medulloblastoma, we analyzed PRC2 subunit expression in SHH medulloblastoma at single cell levels. A recent single cell study of human medulloblastomas (20), which used the Smart-seq protocol that has better sequencing depth, enabled us to examine PRC2 subunit levels in single cells. In all three available SHH medulloblastoma samples representing three SHH medulloblastoma subtypes that occur in infants, children, and adults, respectively, PRC2 subunits displayed heterogeneous levels in cancer-propagating cells that were positive for a CGNP marker ATOH1 (Figure 6E, single cell columns are arranged by the sum of 4 PRC2 subunit levels). In these ATOH1^+^ SHH medulloblastoma cancer propagating cells, PRC2 levels varied from high to undetectable for all 4 subunits. About 10-30% of ATOH1+ cells had no detectable levels of PRC2 subunits in these SHH medulloblastoma samples. PRC2 heterogeneity had no correlation with levels of the SHH medulloblastoma marker GLI2 as shown by similar levels of GLI2 expression throughout the ATOH1^+^ cells regardless of PRC2 levels. Interestingly, higher PRC2 levels in single medulloblastoma cells appeared to correlate with higher levels of the proliferation marker MKI67 (Figure 6E). This result is consistent with an intrinsic oncogenic function of PRC2 in medulloblastoma. PRC2^high^ medulloblastoma cells could have a higher proliferation rate than PRC2^low^ medulloblastoma cells. On the contrary, the presence of the slow-proliferating PRC2^low^ tumor cells suggests that they also contribute to the growth of the tumor. Thus, PRC2 heterogeneity is common in human SHH medulloblastoma. Our mouse genetic experiments and human medulloblastoma analyses together support a model in which PRC2 heterogeneity is oncogenic. In a heterogeneous tumor that contains both PRC2^high^ and PRC2^low^ cells, the PRC2^low^ subclones, despite having a lower growth competence, enhance PRC2^high^ subclone growth through non-cell autonomous mechanisms such as IGF2 signaling. The overall enhanced tumor growth is a result of the cooperation between PRC2^high^ and PRC2^low^ cancer cells (Figure 6F).

### An IGF2/PI3K/AKT pathway mediated non-cell autonomous tumor suppressor function of PRC2 in human medulloblastoma Daoy cells

Using the human SHH medulloblastoma cell line Daoy (60), we further investigated the potential IGF2 mediated non-cell autonomous tumor suppressor function of PRC2. We first confirmed that *IGF2* is repressed by PRC2 in Daoy cells. Treating Daoy cultures with the PRC2 enzymatic inhibitor GSK-126 led to increased *IGF2* expression in a dose-dependent manner as shown by RT-qPCR (Figure 7A). Using the CRISPR-Cas9 method, we generated several *EED^ko^* Daoy clones (Figure 7B). An RNA-seq analysis comparing *EED^wt^* and *EED^ko^* Daoy clones (n=2) also showed the increased expression of oncogenes including *IGF2* and *NOTCH/HES1* (Figure 7C). Interestingly, many EED-reregulated genes encode extracellular factors (Figure 7D), which could support a role of PRC2 in regulating tumor growth via non-cell autonomous mechanisms. Using RT-qPCR, we confirmed the increase of *IGF2* expression in *EED^ko^* Daoy clones compared to the control *EED^wt^* clones (Figure 7E). Our data showed that *EED^ko^* cells have similar or slower growth rates compared to *EED^wt^* controls (Figure 7F). Interestingly, a 4:1 mix of *EED^wt^* and *EED^ko^* cells led to increased growth rates compared to the growth rates of *EED^wt^* and *EED^ko^* cells cultured separately (Figure 7F). In addition, the conditioned media collected from *EED^ko^* cells enhanced *EED^wt^* Daoy cell growth more than did the media collected from the control *EED^wt^* cells (Figure 7G). Consistent with the role of the IGF2/PI3K/AKT pathway in mediating the noncell autonomous function of PRC2 in tumor cell growth, conditioned media from the *EED^ko^* cells caused increased AKT phosphorylation in cultures compared to the addition of *EED^wt^* media (Figure 7H). These results showed that, similar to our observation in mouse SHH medulloblastomas, PRC2 heterogeneity in a human SHH medulloblastoma cell line led to increased growth, possibly through increased IGF2 production from the *EED^ko^* clones.

**Figure 7.**
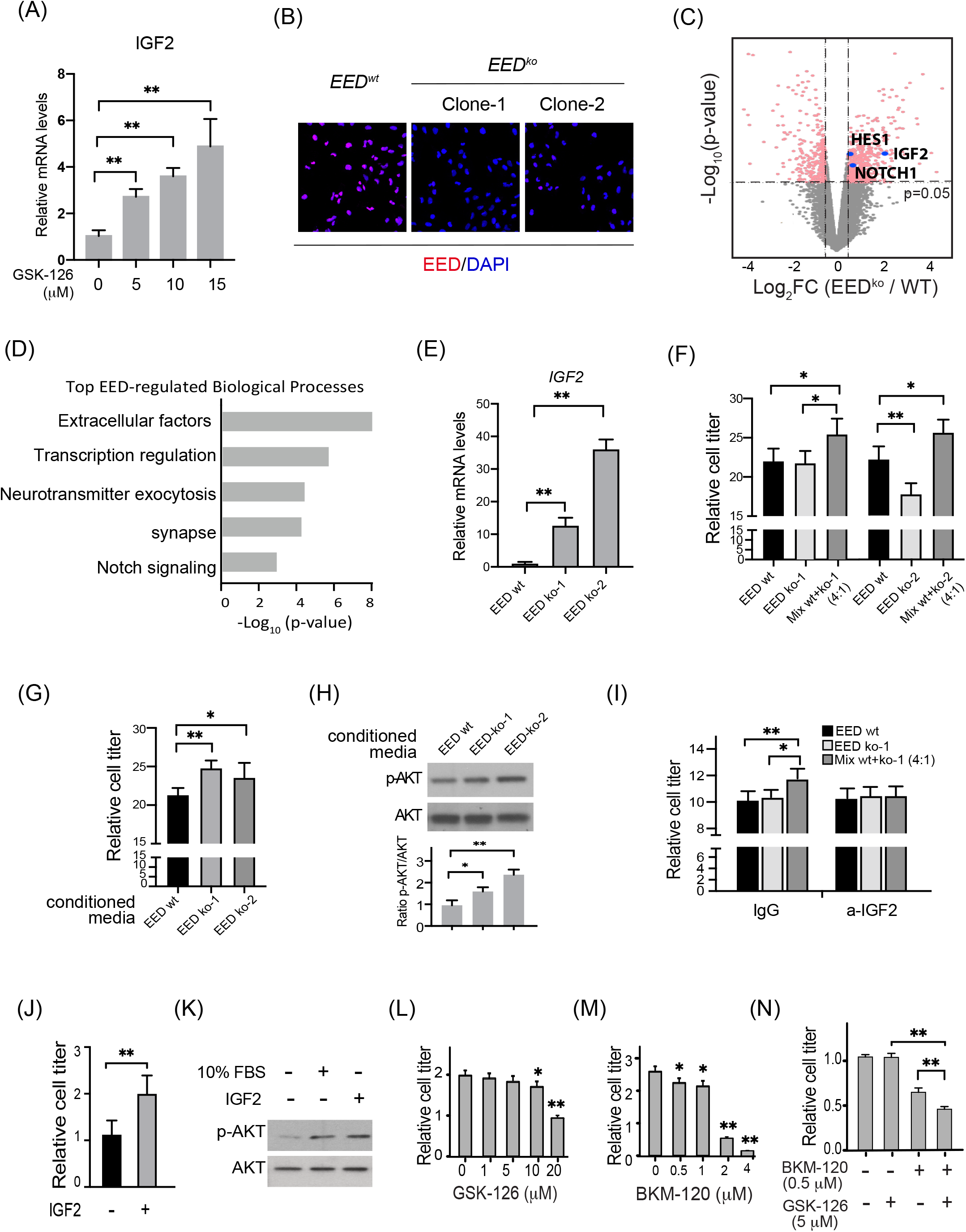
An IGF2/PI3K/AKT pathway mediated non-cell autonomous tumor suppressor function of EED in human medulloblastoma Daoy cells. (A) RT-qPCR of *IGF2* expression in Daoy cell treated with different doses of GSK-126 for 72 hours (n=6). (B) EED staining of *EED^wt^* and *EED^ko^* Daoy cell clones generated by CRISPR-Cas9 system. (C) Volcano plot of RNA-seq data comparing EED^KO^ with *EED^wt^* (n=2). (D) Gene ontology analysis of enriched pathways in up-regulated genes in *EED^KO^* cells. (E) RT-qPCR of *IGF2* expression in *EED^wt^* and *EED^ko^* Daoy cells (n=4). (F) Cell viability assay of *EED^wt^* and *EED^ko^* Daoy cells culturing separately or in a 4:1 mixture (n=5, 72 hours). (G) Cell viability assay of Daoy cells cultured with conditioned media collected from *EED^wt^* or *EED^ko^* cultures (n=5). (H) Western blot showed p-AKT and AKT levels in cells cultured with conditioned media collected from *EED^wt^* or *EED^ko^* cultures. The quantification is shown below (n=3). (I) Cell viability assay of *EED^wt^* or *EED^ko^* or mixed cultures with an IGF2 neutralizing antibody or mouse IgG control (n=5). (J) Cell viability assay of Daoy cells in the absent or present of 200 ng/ml recombinant IGF2 (n=5). (K) Western blot showed p-AKT levels in Daoy cells in different culture conditions. (L-N) Cell viability assay of Daoy cells treated with GSK-126 (L), BKM-120 (M), or in combination (N) for 72 hours at different concentrations (n=5). Student’s t-test, *, p<0.05; **, p<0.01.

To determine whether IGF2 is a key PRC2 target mediating the enhanced growth of Daoy cells with heterogeneous levels of PRC2, we applied neutralizing IGF2 antibodies to the mixed culture of *EED^wt^* and *EED^ko^* cells. The enhanced growth of the mixed culture diminished upon blocking IGF2 signaling (Figure 7I). On the contrary, adding recombinant IGF2 in the media significantly increased growth of Daoy cells (Figure 7J), possibly by increased IGF2/PI3K/AKT pathway activity (Figure 7K). Together, our data suggest a novel oncogenic pathway in SHH medulloblastomas driven by PRC2 heterogeneity through derepressed IGF2 signaling and the downstream PI3K/AKT pathway. Thus, PRC2 inhibition might be able to sensitize tumors to IGF2/PI3K/AKT inhibitors. The PRC2/IGF2/PI3K/AKT pathway could provide effective combination treatment targets for SHH medulloblastoma. Using the Daoy culture system, we tested inhibitors of PRC2 and PI3K separately and in combination. A 72 h treatment with the PRC2 enzymatic inhibitor GSK-126 inhibited Daoy growth, but only at a higher concentration (20 μM) (Figure 7L). The PI3K inhibitor BKM-120 inhibited Daoy cell growth in a dose-dependent manner (Figure 7M). Importantly, a combination treatment using both BKM-120 and GSK-126 at sub-optimal concentrations produced more significant inhibition of Daoy growth than did each individually (Figure 7N). Thus, the oncogenic function of PRC2 heterogeneity and the downstream signaling pathways we discovered may lead to novel treatment strategies for medulloblastoma.

## Discussion

In this study, we report a novel oncogenic function of PRC2 heterogeneity in medulloblastoma development (Figure 6F). We showed that PRC2 heterogeneity is common in human medulloblastoma cancer propagating cells. Using two mouse models and a human cancer cell line, we showed that PRC2 heterogeneity enhances tumor growth by its intrinsic oncogenic function and a novel non-cell autonomous tumor suppressor function. We revealed that the PRC2 repressed IGF2/PI3K/AKT pathway plays a key role in mediating the non-cell autonomous function of PRC2 in enhancing tumor growth. The mechanism we uncovered may shed light on new therapeutic development for medulloblastoma and possibly other cancers.

Tumor growth is driven by intrinsic growth competence of cancer cells and fueled by signals derived from non-cancer cells and other cancer cells in the TME. The understanding of the competitive and cooperative interactions between different cancer subclones could lead to new therapeutic strategies. We revealed a novel oncogenic mechanism arising from PRC2 heterogeneity in cancer subclones. This is supported by several pieces of evidence. Firstly, we generated two mouse medulloblastoma models that contain PRC2^high^ and PRC2^low^ cancer cells using different genetic approaches. Both mosaic models demonstrated significantly enhanced growth compared to homogeneous tumors (Figure 3, 5). Secondly, we generated *EED^wt^* and *EED^ko^* clones of the human medulloblastoma cell line Daoy. Using mixed cultures or conditioned media culture, we showed that *EED^ko^* clones could stimulate the growth of *EED^wt^* clones (Figure 7). Finally, PRC2 heterogeneity is common in human SHH medulloblastoma. The Smart-seq analyses of three SHH medulloblastoma samples showed the presence of PRC2 heterogeneity in ATOH1+ cancer propagating cells in all samples. Importantly, the association between PRC2^high^ cells with a higher expression of the proliferation marker MKI67 demonstrated the different proliferation competence of different subclones (Figure 6D).

We identified IGF2/PI3K/AKT as one key downstream pathway mediating the oncogenic function of PRC2 heterogeneity. *IGF2* is repressed by PRC2 in both mouse and human SHH medulloblastomas (Figure 4, 5C, 6B, 7A, 7E). *IGF2* is highly expressed in SHH medulloblastoma compared to other types of medulloblastomas (Figure 6A). In mouse models, IGF2 has been shown to be increased during CGNP transformation and medulloblastoma initiation and is both necessary and sufficient for tumor progression (54, 59). Downstream of IGF2, high levels of p-AKT were observed in some SHH medulloblastoma patients with poor prognoses (16). We observed increased p-AKT levels in mosaic *Eed* mutant medulloblastomas compared to wild-type medulloblastomas (Figure 4G) or in Daoy cells in response to the conditioned media collected from *EED^ko^* cells or recombinant IGF2 (Figure 7H, 7K). Therefore, the PRC2/IGF2/PI3K/AKT pathway we identified could be a key paracrine/autocrine oncogenic pathway in SHH medulloblastoma progression. Finally, we showed that PRC2 inhibition could sensitize Daoy cells to PI3K inhibitors (Figure 7N).

Besides IGF2, we also identified additional candidate PRC2-repressed oncogenes such as *NOTCH1/HES1* and *LIN28b* (Figure 4D, 7C). Interestingly, both *NOTCH1* and *LIN28b* have been shown to have non-cell autonomous oncogenic functions. Overexpression of NOTCH1 in cerebellum progenitors in the absence of p53 led to the development of SHH type medulloblastoma in mice (61). The oncogenic function appears to be non-cell autonomous since NOTCH1 was only overexpressed in a small number of medulloblastoma tumor cells (61). The NOTCH1/HES1 pathway also plays a non-cell autonomous oncogenic function in small cell lung cancer (10). *LIN28b* is an oncogene that is highly expressed in SHH medulloblastoma (62) and encodes an RNA binding protein that functions through *let-7* miRNA dependent or independent mechanisms. Interestingly, LIN28b also could function non-cell autonomously in development and cancer by promoting IGF2 protein levels (63–65). Thus, these oncogenes may contribute to the oncogenic function of PRC2 heterogeneity. It would be interesting to see whether they function through IGF2-dependent or through independent pathways.

A critical question in tumor heterogeneity and tumor evolution research is the competitive and cooperative relationships between various cancer subclones with different genetic and epigenetic properties. Our study provides an ideal model to study cancer subclone competition and cooperation. In a tumor with subclones with different PRC2 levels, the PRC2^low^ subclones grow more slowly and represent a smaller portion of the heterogenous tumor than do PRC2^high^ subclones. However, PRC2^low^ subclones would not be lost due to their supportive function required for the overall tumor growth. Based on the differential growth competence of the two subclones and the stimulation effects of the PRC2^low^ subclones on the PRC2^high^ subclones, the two subclones may reach a relatively steady ratio (7). The resulting heterogeneous tumors would gain a growth advantage over homogeneous tumors. Although no PRC2 mutation was identified in medulloblastoma, the heterogeneous expression levels of PRC2 in SHH medulloblastoma samples suggest that epigenetic mechanisms suppress PRC2 expression in some cancer cells. It is also possible that mutations of PRC2 subunits could happen in a small number of medulloblastoma cells.

PRC2 plays context-dependent roles in cancer development with both oncogenic and tumor suppressor functions. In medulloblastoma, PRC2 supports cancer cell intrinsic growth competence by repressing CGNP differentiation and cell death. On the other hand, PRC2 also suppresses tumor growth by repressing the expression of secreted oncogenic factors. Together, the pleiotropic functions of PRC2 in cooperative cancer subclones underlie the oncogenic function of PRC2 heterogeneity, which may occur in many other cancer types and may guide the development of new therapeutic strategies.

## Methods

### Mice

The *Atoh1-Cre* mice (46) were provided by Dr. Lin Gan (Rochester University) and Dr. Jane Johnson (University of Texas Southwestern Medical Center). The *Atoh1^+/-^* mice were provided by Dr. Helen Lai (University of Texas Southwestern Medical Center). The *TgAtoh1-Cre* mice (66), the *SmoM2* mice (51), the *CAG-CreER* mice (67), the *Ai9* reporter line (53), and the *Eed^F/F^* mice (47) were purchased from Jackson Laboratory. The *Eed^F/F^* mice contain *loxP* sites flanking exon 3-6 of the *Eed* gene. The NOD/SCID mice were purchased from the University of Texas Southwestern Mouse Breeding Core Facility. All mice are maintained on a mixed genetic background at the University of Texas Southwestern Medical Center Animal Facility. Both males and female mice were used for analyses with no significant differences between them. All procedures were performed in accordance with the IACUC-approved protocols.

### Immunostaining, image processing and BrdU tracing

Paraffin sections of brain or tumor tissues were used for hematoxylin and eosin (H&E) staining as well as immunostaining. For the proliferation assay, BrdU (10 mg/kg; B5002; Sigma-Aldrich) was injected intraperitoneally (adult) or subcutaneously (neonatal) 30 minutes before harvesting. For cell lineage tracing, BrdU (10 mg/kg; B5002; Sigma-Aldrich) was injected subcutaneously in neonatal pups 24 hours before harvesting. The antibodies used for IHC were against Ki67 (eBioscience), BrdU (G3G4, DHSB), GFAP (556330, BD Biosciences), HuC/D (ab184267, Abcam), H3K27me3 (07-449, Millipore), RFP (600-401-379, Rockland), NEUROD2 (ab104430, Abcam), NeuN (ABN78, Millipore), Caspase 3 (9664, Cell Signaling Technology), EED (85322, Cell Signaling Technology) and CRE (15036, Cell Signaling Technology). For IHC staining, biotinylated goat anti-rabbit (BA-1000, Vector Laboratories) or goat anti-mouse (BA-9200, Vector Laboratories) secondary antibodies were used. The counterstain used was either hematoxylin (H-3401, Vector Laboratories) or methyl green (H-3402, Vector Laboratories). For immunofluorescent staining, Alexa 488 conjugated goat anti-rabbit (Thermo Fisher A-11008) and Alexa 594 conjugated goat anti-mouse (Thermo Fisher A-11032) secondary antibodies were used. For the EED and CRE IHC co-stain, three consecutive sections were used to stain EED, CRE, and EED/CRE. The co-stain of EED and CRE was performed following the protocol IHC Multiplexing Guide from VECTOR Laboratories. Bright-field images were acquired using a Hamamatsu Nanozoomer 2.0 HT whole slide scanner at the University of Texas Southwestern Medical Center Whole Brain Microscopy Facility. Fluorescent images were acquired by a confocal system (Zeiss LSM710). Quantifications were performed by counting positive cells from three images per tumor and processed using ImageJ. At least three biological replicates were used in each experiment.

### Tumor transplantation

Medulloblastoma spontaneously grown from *CAG-CreER SmoM2 Eed^F/F^* mice were dissected and dissociated. The tumor cells (5×10^6^) were mixed with Matrigel (BD Biosciences) and injected subcutaneously in the flank regions of NOD/SCID mice. One day after tumor injection, mice were injected intraperitoneally with oil solvent or one or 10 doses of tamoxifen (75 mg/kg) every other day during a 20-day period before being sacrificed for tumor analyses.

### Flow cytometry

Medulloblastoma were dissected and dissociated. Cells were fixed and permeabilized in 4% PFA with 0.1% saponin (Sigma-Aldrich 47036) for 30 min at 4 °C. The primary antibodies used were against EED (85322, Cell Signaling Technology) and CRE (15036, Cell Signaling Technology). The secondary antibody used was anti-rabbit IgG-PE (111-116-144, Jackson Immunoresearch Laboratory). Stained cells were analyzed with a BD FACSCalibur (BD Bioscience). The Flowjo software was used for data analyses.

### Chromatin immunoprecipitation (ChIP)

The ChIP experiments were performed as described previously (30). Dounced tumors were crosslinked with 4% PFA and sonicated into fragments. The antibodies used were against H3K27me3 (07-449, Millipore) and EED (85322, Cell Signaling Technology). The precipitated DNA was purified and subjected to real-time PCR. The graphics shown are representative of experiments performed in triplicate. The experiments were repeated three times.

### RT-PCR, qPCR, and genotyping PCR

The RNA from Daoy cells or tumor tissues was extracted with TRIZOL (Invitrogen). cDNAs were synthesized by reverse transcription using Iscript (Bio-Rad), followed by PCR or quantitative PCR analysis. A Bio-Rad real-time PCR system (C1000 Thermal Cycler) was used for quantitative PCR. *GAPDH* was used to normalize input RNA. The PCR primer sequences were listed in Table S1. Genomic DNA for genotyping was isolated from tails or tumor tissues using a PBND (PCR buffer with nonionic detergents) preparation. A Bio-Rad C1000 Thermal Cycler was used for PCR. The PCR primer sequences were listed in Table S1.

### RNA-seq analyses

For RNA-seq, total RNAs were extracted, followed by library preparation, using the Illumina RNA-Seq Preparation Kit and sequencing on a HiSeq 2500 sequencer at the University of Texas Southwestern Sequencing Core Facility. The expression levels of genes were quantified by a Kallisto program (68), and the differentially expressed genes were detected by an EdgeR program (69). Genes with a count-per-million (CPM) of less than 1 in more than 2 samples were excluded. The differentially expressed genes with a fold change larger than 2 and an FDR<0.05 were selected as EED-regulated genes. Gene ontology analysis was performed using DAVID tools (http://david.abcc.ncifcrf.gov/).

### Analyses of human medulloblastoma expression data

The gene expression data from subtype classified human medulloblastoma were extracted from previous studies (16, 58). The Kaplan-Meier method was used to compare the survival probability of patients with different tumor expressions of PRC2 subunits (58). Single cell Smart-seq expression data were extracted from a previous study (20). The expression of candidate genes in all ATOH1+ cells were analyzed.

### Cell cultures

The medulloblastoma cell line Daoy was purchased from ATCC and was cultured as suggested by the supplier. Conditioned media were collected from Daoy cultures 24 hours after changing to the serum free medium and were used to culture *EED^wt^* Daoy cells for 72 hours before measuring cell viability. For co-culture experiments, *EED^wt^, EED^ko^*, or mixed (4 *EED^wt^*:1 *EED^ko^*) cells were seeded in 96-well plate at 5,000 cells per well. After culturing for 24 hours in complete media (containing 10% FBS), culture media were changed to serum free media or serum free media with 2 ug/ml IGF2 antibody (AF-292, R&D Systems) for another 72 hours, or 200 ng/ml recombinant human IGF2 (292-G2, R&D Systems) for 24 hours before measuring cell viability. Relative cell titer was determined using the CellTiter-Glo^®^ Luminescent Cell Viability Assay (Promega, G7572) according to the manufacture’s instruction. For inhibitor experiments, GSK-126 (S7061, Selleckchem), BKM-120 (S2247, Selleckchem), or the two in combination were added to culture media for 72 hours before measuring cell viability.

### Lentivirus preparation and infection

For packaging the lentivirus, CRISPR-cas9-based guide RNA (gRNA) targeting human *EED* (GATCATAACCAACCATTGTT) was cloned in LentiCRISPR v2 (Addgene 52961), which was mixed with psPAX2 and pMD2.G (Addgene) and transfected into HEK 293T cells using Polyjet (Signagen). Supernatant containing the viruses was collected 48-72 hours after transfection and used for subsequent infection. Daoy cells were infected with lentiviral supernatant containing 8 ug/ml polybrene for 24 hours. In order to select *EED^ko^* Daoy cells, puromycin (1 μg/ml) was added three days after initial infection for one week. Single cell clones were selected by limiting dilution. EED deletion was confirmed by immunofluorescent staining.

### Immunoblotting

Daoy cells were lysed in RIPA buffer (50 mM Tris, pH 8, 250 mM NaCl, 0.05% SDS, 0.5% DOC, 1% NP-40) in the presence of phosphatase inhibitors. The cell lysates were separated on 12% SDS-PAGE gels. The antibodies used were against AKT (sc-8312, Santa Cruz Biotechnology) and p-AKT (4060, Cell Signaling Technology).

### Statistical analysis

The data are expressed as means ± s.d. All experiments were repeated for 3 times unless otherwise indicated. Statistical analysis was performed by either an ANOVA post hoc t-test for multiple comparisons or a two-tailed unpaired Student’s t-test. For the morbidity studies, the Kaplan-Meier method was used to plot the survival curve, and the log-rank test was used for statistical analyses. A p-value < 0.05 was considered significant.

## ACKNOWLEDGEMENT

We are grateful to Drs. Jane Johnson and Lin Gan for providing the Atoh1-Cre mice and Dr. Helen Lai for providing the Atoh1+/- mice. We thank Dr. Zaili Luo for assisting the human medulloblastoma expression data analyses. We thank Dr. Xiaoye Liu and Mr. Huaxia Dong for technical help, and Michael Zhan for reading the manuscript. This work was supported by grants from NIH (R01NS096068 and R21NS1045968 to J.W.).

## CONTRIBUTIONS

J.W. and J.Y. designed the experiments and analyzed the results. J.Y., X.S, and X.Z performed the experiments and collected the data. R. L. provided human medulloblastoma expression and prognosis data. Z.X. performed bioinformatics analyses. J.Y. and J.W. wrote the manuscript with the help from all authors.

## CONFLICT OF INTEREST

The authors declare no competing interest.

## Notes

### Competing Interest Statement

The authors have declared no competing interest.

