## Supplemental figures and table for "A Novel Oncogenic Function of PRC2 Heterogeneity in Medulloblastoma"

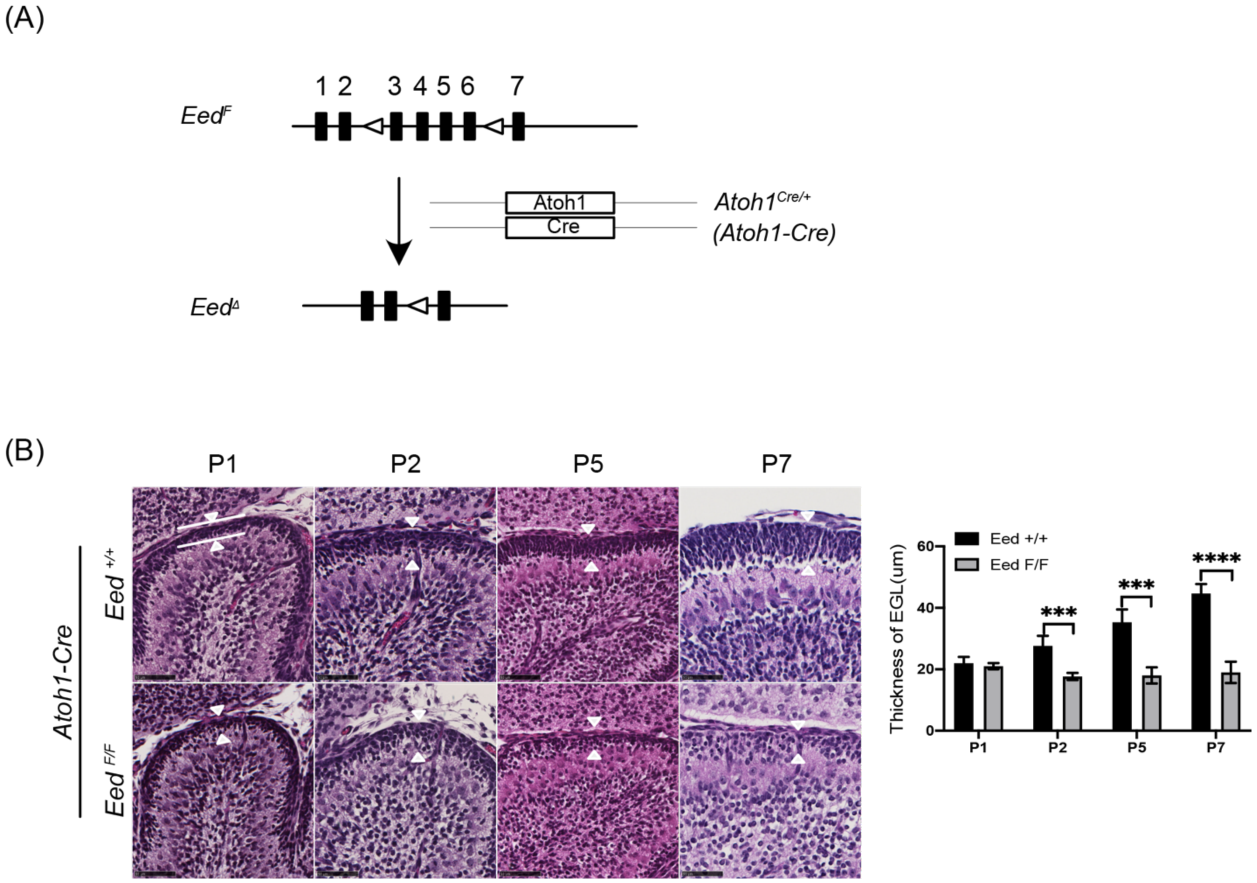


**Figure S1. Atoh1-Cre mediated *Eed* deletion led to CGNP depletion.**

(A) The genetic strategy for deletion of floxed *Eed* (*Eed*$^{F}$) allele in CGNPs with the knockin knockout *Atoh1-Cre (Atoh1*$^{Cre/+}$*)*. LoxP sites flanking the *Eed* exons are shown as empty triangles. (B) H&E staining of sagittal sections of cerebella from P1, P2, P5, and P7 *Atoh1-Cre Eed*$^{+/+}$ and *Atoh1-Cre Eed*$^{F/F}$ mice. Quantification of the thickness of EGL is shown on the right (n=3). Student's t-test, ***, p<0.001; ****, p<0.0001. **Related to Figure 1.**

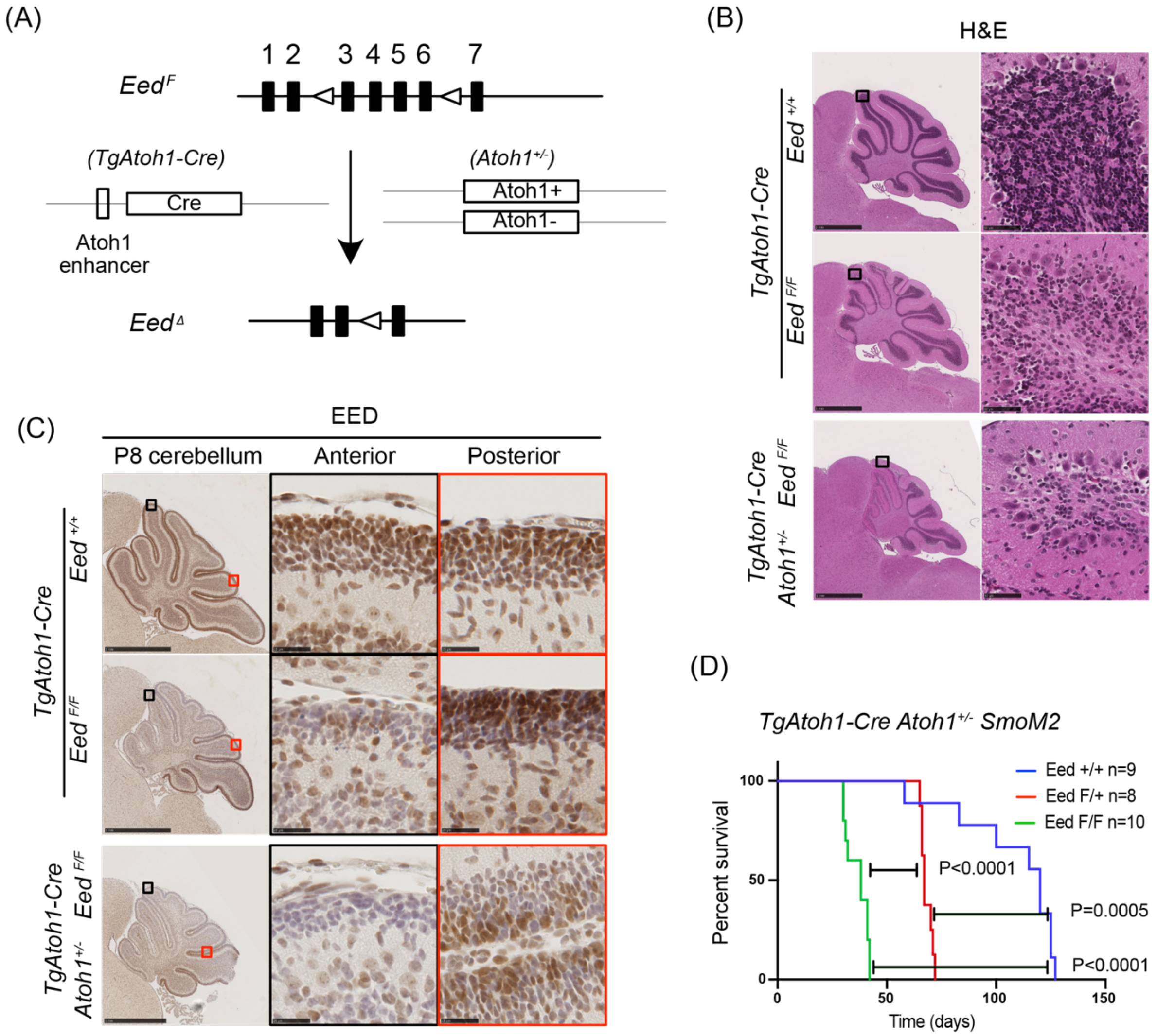


**Figure S2. The effects of *Atoh1* heterozygosity on the phenotypes of *TgAtoh1-Cre* mediated *Eed* deletion.**

(A) The genetic strategy for deletion of floxed *Eed* (*Eed*$^{F}$) allele in CGNPs with *TgAtoh1-Cre*. LoxP sites flanking *Eed* exons are shown as empty triangles. (B) H&E staining of sagittal sections of cerebella from P28 *TgAtoh1-Cre Eed*$^{+/+}$, *TgAtoh1-Cre Eed*$^{F/F}$, and *TgAtoh1-Cre* Atoh1$^{+/-}$ *Eed*$^{F/F}$ mice. Pictures on the right are boxed areas in the pictures on the left. (C) EED staining of sagittal sections of cerebella from P8 *TgAtoh1-Cre Eed*$^{+/+}$, *TgAtoh1-Cre Eed*$^{F/F}$, and *TgAtoh1-Cre* Atoh1$^{+/-}$ *Eed*$^{F/F}$ mice. Pictures on the right are boxed areas in the pictures on the left with corresponding colors. (D) Survival curves of *TgAtoh1-Cre SmoM2* mice with different *Eed* genotypes and with or without *Atoh1* heterozygosity. **Related to Figure 2 and Figure 3.**

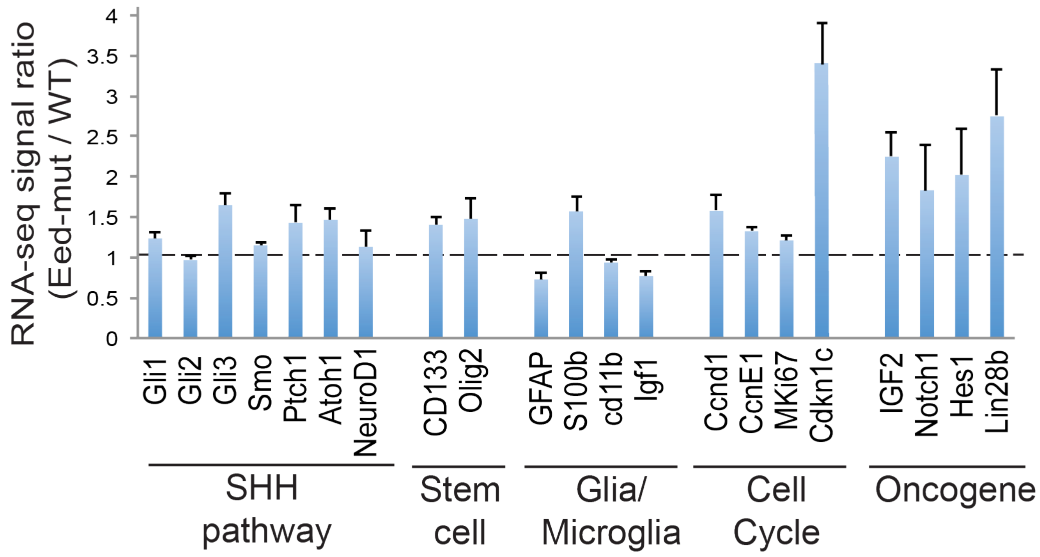


**Figure S3.** RNA-seq signal ratios of key SHH medulloblastoma genes between *TgAtoh1-Cre SmoM2* $Eed^{F/F}$ (Eed-mut) and *TgAtoh1-Cre SmoM2* $Eed^{+/+}$ (WT) medulloblastomas. **Related to Figure 4.**

### Table S1. Sequences of PCR primers

| **RT-PCR Primers** | Sequence | |
|---|---|---|
| Gene (Mouse) | Forward | Reverse |
| Igf2 | GTGCTGCATCGCTGCTTAC | ACGTCCCTCTCGGACTTGG |
| Notch1 | CCCTTGCTCTGCCTAACGC | GGAGTCCTGGCATCGTTGG |
| GAPDH | GTGGTGAAGCAGGCATCTGA | GCCATGTAGGCCATGAGGTC |
| Gene (Human) | | |
| IGF2 | CTTGGACTTTGAGTCAAATTGG | GGGTCGTGCCAATTACATTTC |
| GAPDH | CCACCCATGGCAAATTCC | TGGGATTTCCATTGATGACAAG |
| **ChIP primers** | | |
| Gene (Mouse) | | |
| Igf2 | AACTGAGCACAGGTCCTGCC | CGAGGTCCCATGTCATGTTTC |
| Hoxa7 | CTCTTCTGTTTCCCATCCTGGT | GGCAATATCCGGGATCCACT |
| **Genotyping primers (mouse)** | | |
| $Eed^{F}$ | GGGACGTGCTGACATTTTCT | CTTGGGTGGTTTGGCTAAGA |
| $Eed^{\Delta}$ | GGGACGTGCTGACATTTTCT | CTTTGGCATTTATGATAAAGC |
| Cre | TCGATGCAACGAGTGATGAG | TCCATGAGTGAACGAACCTG |
